# Rapid detection of SARS-CoV-2 and other respiratory viruses by using LAMP method with Nanopore Flongle workflow

**DOI:** 10.1101/2020.06.03.131474

**Authors:** Jingjing Li, Weipeng Quan, Shuge Yan, Shuangju Wu, Jianhu Qin, Tingting Yang, Fan Liang, Depeng Wang, Yu Liang

## Abstract

The ongoing novel coronavirus (COVID-19) outbreak as a global public health emergency infected by SARC-CoV-2 has caused devastating loss around the world. Currently, a lot of diagnosis methods have been used to detect the infection. The nucleic acid (NA) testing is reported to be the clinical standard for COVID-19 infection. Evidence shows that a faster and more convenient method to detect in the early phase will control the spreading of SARS-CoV-2. Here, we propose a method to detect SARC-Cov-2 infection within two hours combined with Loop-mediated Isothermal Amplification (LAMP) reaction and nanopore Flongle workflow. In this approach, RNA reverse transcription and nucleic acid amplification reaction with one step in 30 minutes at 60-65°C constant temperature environment, nanopore Flongle rapidly adapter ligated within 10 minutes. Flongle flow cell sequencing and analysis in real-time. This method described here has the advantages of rapid amplification, convenient operation and real-time detection which is the most important for rapid and reliable clinical diagnosis of COVID-19. Moreover, this approach not only can be used for SARS-CoV-2 detection but also can be extended to other respiratory viruses and pathogens.

## Introduction

Until April 2020, the outbreaking COVID-19 around the world caused by SARS-CoV-2 have resulted in almost 3,000,000 infection cases and more than 200,000 deaths. Early studies show that COVID-19 have an incubation period time when you catch a virus until your symptoms start. During the incubation period time SARS-CoV-2 can also occur Human-To-Human COVID-19 transmission. At this moment, there is no effective vaccine or drugs for the COVID-19. Therefore, rapidly and conveniently diagnosis of the patients in the early phase was supposed to the most effectively way to control the COVID-19 epidemic.

Here, we use nanopore Flongle workflow combined with LAMP reaction to propose a faster and more convenient method to detect SARS-CoV-2 and other respiratory viruses in two hours. LAMP amplification is a rapid, convenient, highly sensitive and specific technique for clinical samples. Amplification reaction can be done less than 30 minutes in a friendly environment. Nanopore Flongle is designed to be the quickest, most accessible and cost-efficient for real-time sequencing. We use the nanopore Flongle workflow for genome sequencing and analysis after sequencing 30 minutes for SARS-CoV-2 identification.

This study presents a LAMP based method combined with nanopore Flongle rapid realtime sequencing workflow to detect COVID-19 as low as 3.25×10^2 copies/mL of SARS-CoV-2 in both laboratory and wild-caught environment. It takes less then two hours to diagnose the COVID-19 from the RNA isolate. In additional, it does not require highly trained people and does not involve expensive and sophisticated equipment for amplification. In summary, this approach can be used at the point of care by field and local personnel for the rapid diagnosis of SARS-CoV-2 as well as the investigation of outbreaks of other respiratory diseases in the future.

## Methods

We propose a fast and efficient method (Figure 1) for the detection of COVID-19 infection and other respiratory viruses. The method described here can detect SARS-CoV-2 and Influenza B virus within two hours at a low copy to 3.25×10^2 copies/mL in a friendly environment.

**Figure 1.**
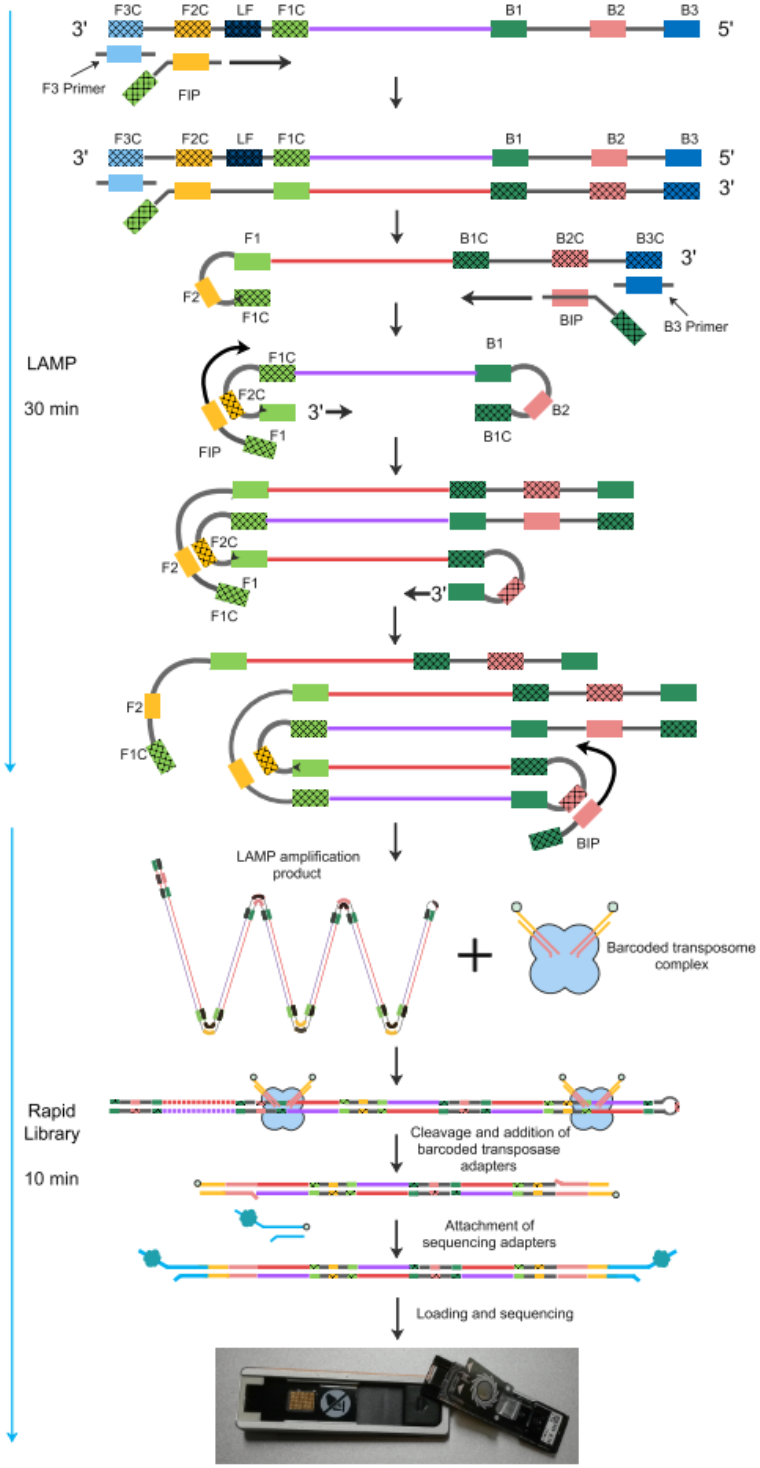
The detection method for COVID-19 combined LAMP with nanopore Flongle

### LAMP Primer Design

The species-specific genes (Suppl. Table 1) were chosen as target genes for SARS-CoV-2 and Influenza B virus. All the target genes were synthesized and then cloned into a pGEM-T easy vector by TsingKe (Wuhan, China). Primers for LAMP reaction were synthesized by Invitrogen biotech (Shanghai, China). Each primer (Suppl. Figure 1) contain 6 oligonucleotides respectively forward inner primer (FIP), backward inner primer (BIP), forward outer primer (F3), backward outer primer (B3), loop forward primer (LF), loop backward primer (LB). FIP, BIP, F3, B3, these 4 primers involve 6 independent sequences of the target region. All primers were aligned to the published SARS-CoV-2 sequences and others respiratory using BLAST. Results show the primers chosen were specific.

### LAMP Reaction

Using Bst 2.0 DNA polymerase (NEB, M0537) by self-recycling strand displacement to synthesize DNA. Since the plasmids were used as amplification template, the reverse transcription (RT) can be omitted (approximately 15 minutes). The plasmids of SARS-CoV-2, Influenza B virus target genes were equimolar pooling into one tube and diluted at concentration of 3.25×10^4, 3.25×10^3, 1.1×10^3, 6.5×10^2, 3.25×10^2 copies/mL. All the dilution gradient and pure water (negative control) were amplified by LAMP. A LAMP reaction contains 2.5 μL 10X Isothermal Amplification Buffer, 1.5μL MgSO_4_ (100 mM), 3.5 μL dNTP Mix (10 mM each), 1 μL Bst 2.0 DNA Polymerase (8,000 U/ml), primers (final concentration: inner primers 1.6 μM, outer primers 0.2 μM, loop primers 0.4 μM), 10μL target DNA sample, adjust the volume to 25 μL with nuclease-free water. The reaction was run in a thermal cycler (65° C, 30 minutes). The gel electrophoresis image (Suppl. Figure 2) of amplification was smear band, which like staircases, and the longest fragment was up to 10 kb. The reaction solution was purified by 35μL AMPure XP beads (Beckman, A63881), and quantified with Qubit^®^ dsDNA HS Assay Kit (Invitrogen, Q32854).

### Nanopore Sequencing and Basecalling

200 ng input 3.25×10^5 copies/mL amplification products was used to construct ONT rapid sequencing library (Oxford Nanopore, SQK-RBK004) according to the manufacturer’s instructions. 10μL total reaction mixture containing 7.5μL of purified amplification DNA and water, 2.5μL Fragmentation mix (FRA) were added together to the reaction tube. Mix well and incubate the reaction tube at 30°C for 1 minute and then at 80°C for 1 minute. Briefly put the tube on ice to cool it down. The FRA is a barcoded transposome complex, which can cleave randomly DNA and add barcoded transposase adapters to the cleavage sites. 1μL Rapid Adapter (RAP) add to 10 μL previous reaction to attach sequencing adapter. Mix well, and incubate the reaction for 5 minutes at room temperature. Then the nanopore library was loaded into Flongle flowcell for real-time sequencing. The sequencing data were processed as describe for pathogen identification after running 30 minutes.

To test the limit of detection, the amplification products of dilution gradient 3.25×10^4, 3.25×10^3, 1.1 × 10^3, 6.5 × 10^2, 3.25×10^2 copies/mL and negative control total 12 samples were constructed another barcoding library (Oxford Nanopore, SQK-RBK004) as described above and sequenced using a PromethION flowcell to achieve more data.

At the same time of real-time sequencing basecalling is doing in the nanopore platform with high accuracy mode by Guppy (version 3.2.10). Quality control was processed by MinKNOW (version 3.6.5) software pipeline in the machine with reads quality value score cutoff 7.

### Detection Method

The reads after quality control called passed reads generated by the nanopore platform was processed in real-time. We use qcat (version 1.1.0) to demultiplex the barcoding samples. Statistical analysis of nanopore long reads sequencing data about reads length distribution, reads count, reads score was processed with an in-house pipeline. We align nanopore long reads to virus genome database with minimap2 (version 2.1) and use samtools (version 1.9) to demonstrate the coverage and depth. We confirm the sample is positive or not for the virus by count the aligned reads ratio, aligned reads count, aligned reads identity.

## Result

### Study design for SARS-CoV-2 detection

The study design (Figure 2) for SARS-CoV-2 detection is based on LAMP rapid amplification of specific genes and sequenced by nanopore Flongle workflow. We use standard plasmids of orf1ab gene, N gene from SARS-CoV-2, plasmids of HA gene, M gene from influenza B virus as a mixed sample. Furthermore, homo sapiens GAPDH gene was added to the mixed genes as one of the reaction controls. The amplified products of those target genes were ligated by nanopore Flongle rapidly adapter, followed by sequencing on nanopore Flongle workflow. For the detection sensitivity of SARS-CoV-2 identification by LAMP reaction and different reaction times, we diluted the mixed sample to 3.25×10^4, 3.25 ×10^3, 1.1×10^3, 6.5×10^2, 3.25×10^2 copies/mL and with LAMP reaction 20 minutes and 40 minutes. Total 11 dilution samples and a negative control (pure water) were prepared for LAMP PCR amplification. In order to generate more data and improve detection sensitivity, we barcoding the samples and pooling together for nanopore PromethION platform. Compared with qPCR testing our method will output nucleic acid sequence from target genes if the sample is positive. So, we can identify whether the sample is the positive or negative by aligned reads ratio, aligned reads count, aligned coverage and identity.

**Figure 2.**
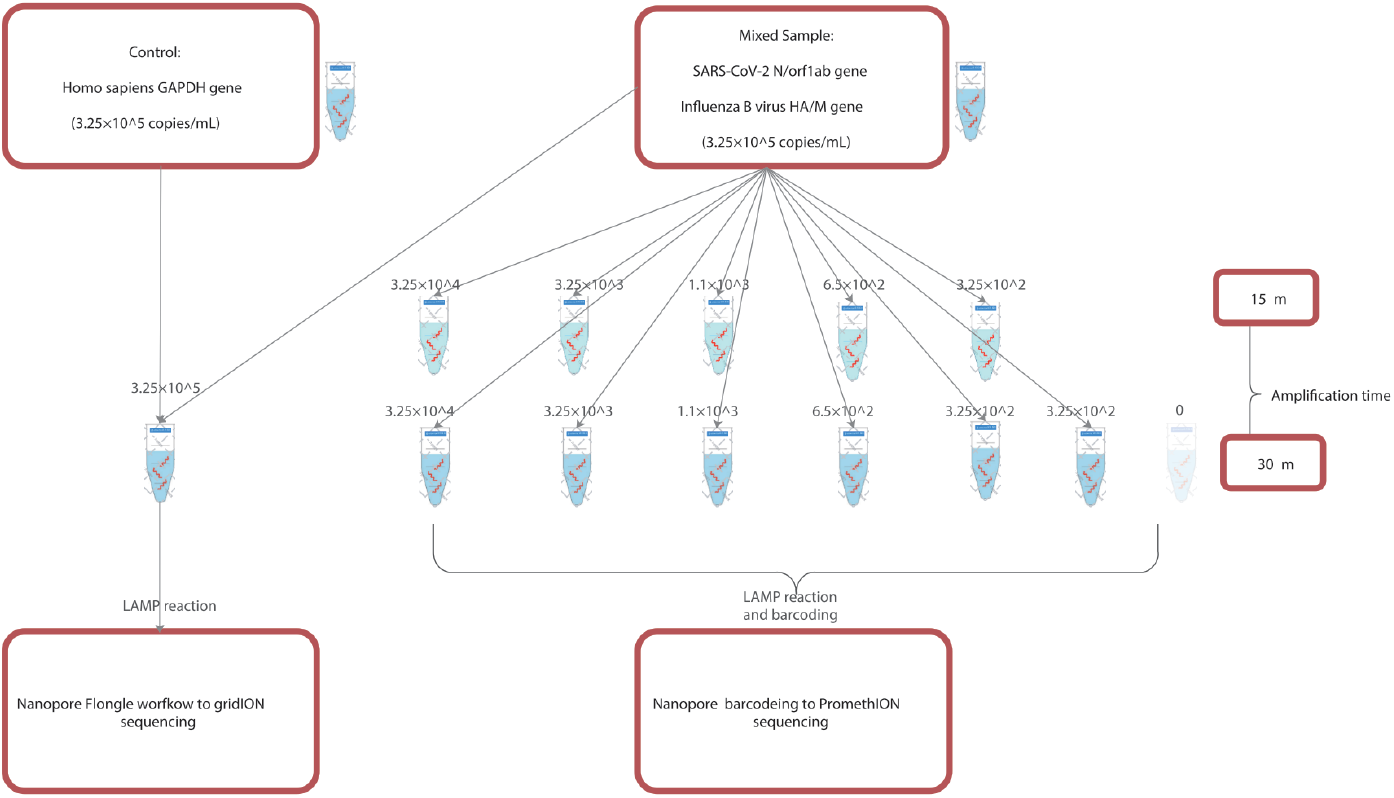
Study design for SARS-CoV-2 detection

To avoid amplification failure and improve the sensitivity of detection. For each microbe we choose two genes as target gene to design primers and dilute the mixed samples to different gradients. We choose SARS-CoV-2 (NC_045512.2), orf1ab gene region (genome 16,500 bp-18,000 bp), N gene region (genome 1 bp-1260 bp) to design LAMP primers. For influenza B virus is HA gene (MT040795), M gene (MT040797) as target gene.

After amplification, we use nanopore Flongle workflow to generate sequences data in real-time. This sequencing model allowed us to align the reads to the genome and process the data in real-time, so we can identify the COVID-19 infection after sequencing in few minutes.

### LAMP Primer design and reaction for SARS-CoV-2

In order to detect the SARS-CoV-2, the species-specific genes orf1ab and N gene were chosen as target genes after aligned to other HCOVID (Human Coronavirus) such as MERS, SARS, OC43, HKU1 that happened to human. LAMP primers were designed by using NCBI Primer-BLAST. The primers were aligned to other coronaviruses and published SARS-CoV-2 sequences in genebank by BLAST alignment on line in NCBI. The aligned results showed the primers we chosen were species-specific. The Influenza B virus primers take the same approach HA gene and M gene were chosen as target gene.

### Nanopore platform sequencing and quality control

A nanopore Flongle flowcell sequencing library was prepared using 200 ng of DNA amplification product (amplification 30 minutes) as input to SQK-RBK004 kit (Oxford Nanopore Technology, UK). This library was sequenced on nanopore GridION device with running 30 minutes, 1 hour, 2 hours, 12 hours for real-time analysis (Suppl. Figure 3A). Nanopore reads were base called using Guppy (version 3.2.10) at a high accuracy mode. Output fastq files were quality controlled by filtering reads quality value 7 called passed reads (Suppl. Figure 3B). For running 30 minutes we got more then 2000 passed reads to detect COVID-19, the max read length is 3038 bp, average reads length is 481 bp, mean reads quality score is 15. After 12 hours finished sequencing total generated 27000 reads, the reads length and quality have the same performance when sequencing 30 minutes (Suppl. Table 2).

Multiplex sequencing libraries after amplification reaction were constructed using 4.4 ng – 200 ng of DNA (amplification 15 minutes to 30 minutes) from up to 12 samples (Table 1) as input to SAK-RBK004 kit and barcoded individually. This library was sequenced on the PromethION device with running 24 hours. After base calling and based quality control, we demultiplex the barcoding libraries by using qcat (version 1.1.0) with min-score (Suppl. Figure 4) 80. The nanopore PromethION is a high-throughput sequencer compare with Nanopore Flongle, we take more stringent barcode score to 80 when using qcat to improve demultiplexed accuracy. The negative control (amplification 30 minutes) after demultiplexed generate 209 reads, probably caused by nanopore barcode demultiplexed error. Compared with other positive samples about reads count the order of magnitude is very small.

**Table 1.**
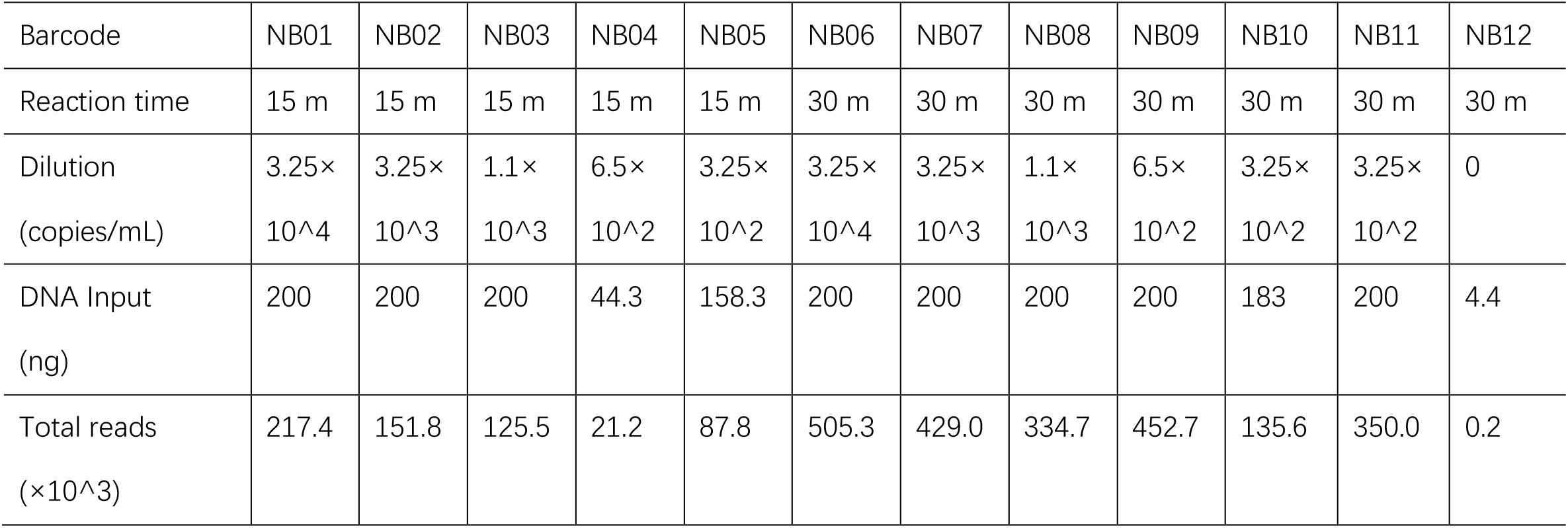
The barcoding samples about different dilution gradients and reaction time

### SARS-CoV-2 detection using nanopore Flongle

In order to achieve a rapid detection of COVID-19 infection, the sequenced data were analysed in real-time. We compare the results by data sequenced time line 30 minutes, 1 hour, 2 hours, 12 hours. Minimap2 (version 2.1) was used to map the nanopore long reads against the reference genome to count the detected reads count with aligning identity 85%. For different running time more than 90% (Figure 3A) of the sequenced reads were aligned to reference genome and have a high identity against the reference. Then the detected reads (Table 2) of each time line was assigned to each target gene (Figure 3B). Aligned reads and assigned reads show uniform performance to target species and target genes (Suppl. Figure 5).

**Figure 3.**
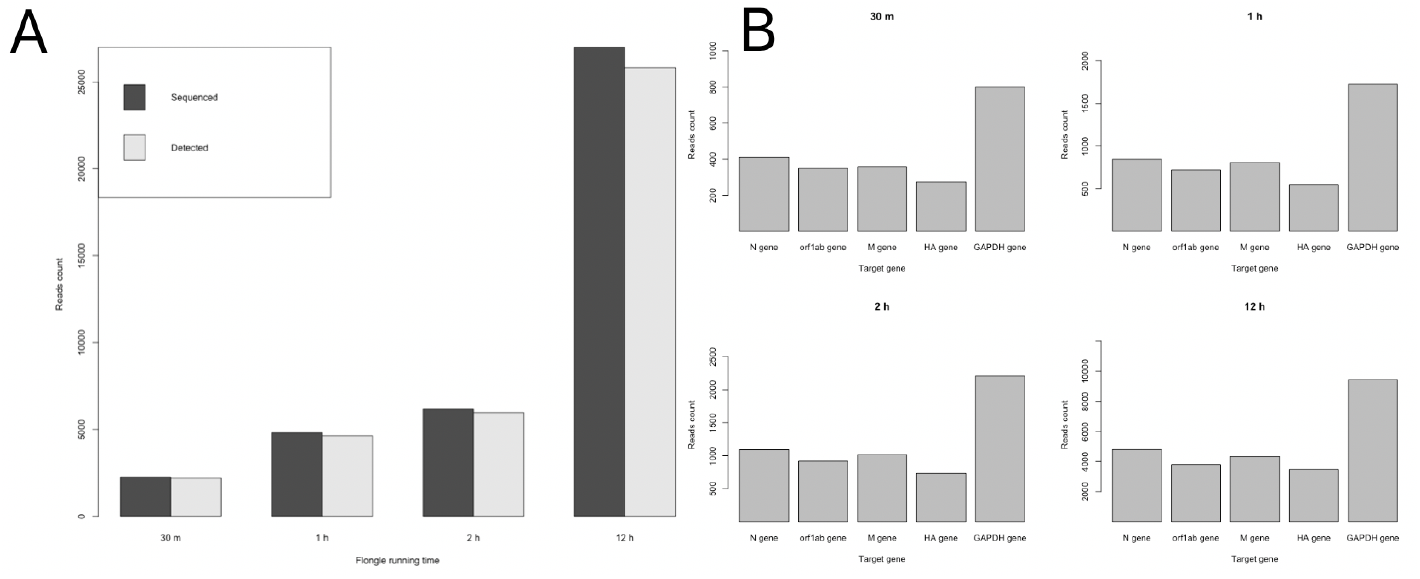
Nanopore Flongle different running time SARS-CoV-2 detection (A) Flongle different running time sequenced and detected reads numbers to reference; (B) Flongle different running time aligned reads assign to different target gene number

**Table 2.** Nanopore Flongle detected reads assigned to target gene

| Species | Target | Reads #(30m) | Reads #(1h) | Reads #(2h) | Reads #(12h) |
| --- | --- | --- | --- | --- | --- |
| SARS-CoV-2 | N gene | 409 | 845 | 1089 | 4786 |
| SARS-CoV-2 | orf1ab gene | 348 | 719 | 916 | 3803 |
| Influenza B Virus | M gene | 359 | 801 | 1013 | 4336 |
| Influenza B Virus | HA gene | 274 | 551 | 732 | 3473 |
| Homo sapiens | GAPDH gene | 801 | 1724 | 2207 | 9421 |

Compared with sequencing 12 hours, sequencing 30 minutes output more than 2000 nanopore reads used to SARS-CoV-2 detection showed the same performance and results. So, we can identify SARS-CoV-2 infection by sequencing few minutes with this method.

### Sensitivity of SARS-CoV-2 identification by nanopore platform

To confirm the detection sensitivity of SARS-CoV-2 by nanopore platform, we use the mixed sample diluted to different gradients (from 0-3.25 ×10^4 genome copies/mL) with different reaction time from 15 minutes to 30 minutes together a negative control total 12 samples. After amplification (Table 1) all the samples were barcoded and loaded to a nanopore PromethION chip sequencing 24 hours.

Barcode after demultiplexing positive samples (Figure 4) from different gradients and reactions generate reads range from 21.2×10^3 to 505.3 ×10^3 reads. For positive samples the detected reads of all sequenced reads ratio approach to 90%. Negative control as an experimental control which should have no reads output, but we got 14 reads among 209 reads aligned to reference about 7% percent. This may be caused by demultiplexed error.

**Figure 4.**
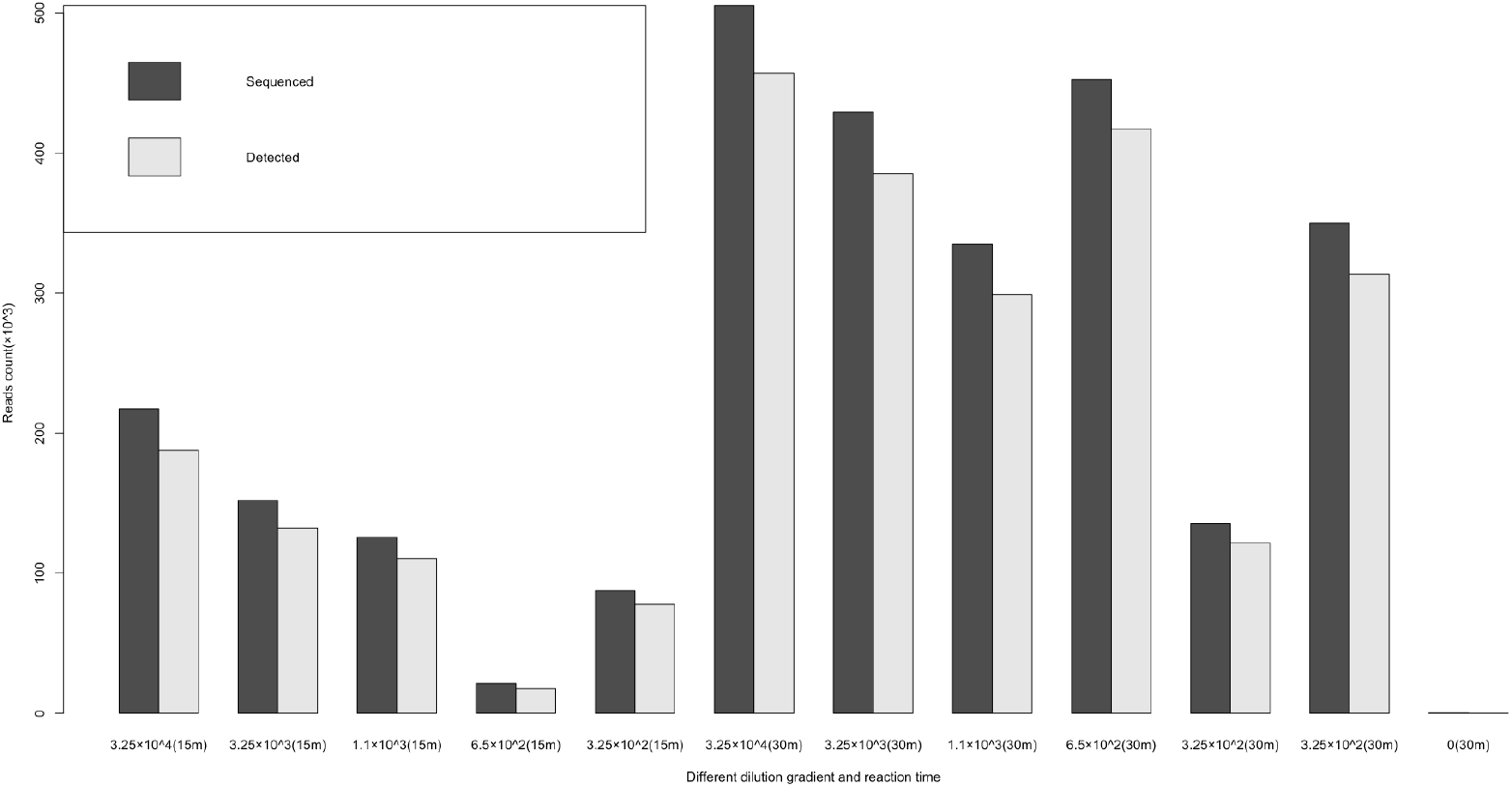
Nanopore demultiplexed and detected reads number of each sample

## Discussion

The outbreaking COVID-19 infection by SARS-CoV-2 around the world has caused devastating loss of economics and thousands of people died. Rapid, accurate and convenient diagnosis for COVID-19 infection and then take a strict restrictive isolation measures is the best way to control COVID-19 threating public health. Here, we proposed a new method to detect SARS-CoV-2 and other respiratory viruses by using LAMP amplification and then incorporated it into a nanopore Flongle sequencing workflow to virus species identification within two hours from sample receipt.

Our workflow for COVID-19 and other respiratory samples detection includes DNA depletion, microbial DNA extraction, target gene amplification, nanopore Flongle sequencing, data analysis. We design primers for detecting orf1ab gene, N gene of SARS-CoV-2 and HA gene, M gene of influenza B virus, GAPDH gene of Homo sapiens as control. The LAMP can complete target nucleic acid amplification in 30 minutes at 60-65°C constant temperature environment. Furthermore, products after LAMP reaction were loaded into nanopore Flongle adpater to sequence and analysis in real-time. To confirm the LAMP have a highly sensitivity, we dilute the samples to 3.25×10^4, 3.25×10^3, 1.1×10^3, 6.5×10^2, 3.25×10^2 copies/mL, results show that for SARS-CoV-2 N gene can be detected after diluted to 10^1 copies/mL.

within two hours have a high sensitivity at 3.25×10^2 copies/mL. As we known LAMP and RT-LAMP have been established for use as a highly sensitive methods for pathogen detection also contain SARS in 2003. We combine LAMP and nanopore Flongle workflow to detect SARS-CoV-2 in two hours.

In conclusion, we report a rapid, accurate and convenient method for COVID-19 infection detection by using LAMP amplification with nanopore Flongle workflow. The LAMP reaction has been demonstrated to be a simple, fast, and highly sensitive method for sequence specific viral nucleic acid detection. The nanopore Flongle workflow was designed to offer rapid, low cost and on demand sequencing in real time. By additional barcoded design on LAMP and individual samples, this method can be scaled to detect millions of samples one day by using Nanopore GridION or PromethION platform. Moreover, this method for COVID-19 rapid detection can also extend to other pathogens by additional primer design.

## Supporting information

Suppl. Nanopore Flongle

