## Supplementary material for "Rapid detection of SARS-CoV-2 and other respiratory viruses by using LAMP method with Nanopore Flongle workflow": Suppl. Nanopore Flongle

[1]GrandOmics Diagnostics, Wuhan, Hubei, 430000 China.
[2]GrandOmics Biosciences, Beijing, 102200 China.
[3]Grandomics Biosciences, Wuhan, Hubei, 430000 China.

[*]These authors contributed equally.
[#]Yu Liang:; Depeng Wang:.

### Supplemental materials figure and table

Suppl. Table 1 Target genes and primers for Lamp amplification

| Species | Target gene | Primer | Sequence |
|---|---|---|---|
| SARS-CoV-2 | N gene | LF | TGGACTGAGATCTTTCATTTTACCG |
| | | LB | ACTGAGGGAGCCTTGAATACA |
| | | F3 | TGGCTACTACCGAAGAGCT |
| | | B3 | TGCAGCATTGTTAGCAGGAT |
| | | FIP | TCTGGCCCAGTTCCTAGGTAGTGACGAATTCGTGGTGGTGA |
| | | BIP | AGACGGCATCATATGGGTTGCAGCGGGTGCCAATGTGATC |
| | orf1ab gene | LF | ACATAGTGCTTAGCACGTAATC |
| | | LB | GGGCACACTAGAACCAGAATATTT |
| | | F3 | TCTTTGATGAAATTTCAATGGC |
| | | B3 | GTTCCGAGGAACATGTCT |
| | | FIP | GAGCAGGGTCGCCAATGTACTTTGAGTGTTGTCAATGCCA |
| | | BIP | CACCACGCACATTGCTAACTAAGGACCTATAGTTTTCATAAGTCTAC |
| Influenza B virus | HA gene | LF | TGCCGCTTTGTGGTAGTCC |
| | | LB | TGTGTTGTTGCCTCAAAAGGT |
| | | F3 | CCACACATTATGTTTCTCAGAT |
| | | B3 | ACCAATTAAAGGCAATGACC |
| | | FIP | CTGCACCATGTAATCAACAACAATAGATCAAACAGAAGACGGA |
| | | BIP | AGGAACAATTGTCTATCAAAGGGGTTACTTTGCTCCTGCCAT |
| | M gene | LF | GCACAAAGCACAGAGCGT |
| | | LB | AGGAGTGAGACGGGAAATGC |
| | | F3 | TGTACCTGAATCCTGGAAAT |
| | | B3 | TTTTGGACGTCTTCTCCTT |
| | | FIP | ATGAGCTCTGTGTGAATGTGATGATTCAATGCAAGTAAAACTAGGA |
| | | BIP | AGAGCAGCGAGATCCTCAGTCATTGTTTTTGCTGTGTTCAT |
| Homo sapiens | GAPDH gene | LF | CGCCCCACTTGATTTTGGAG |
| | | LB | AGGCTGGGGCTCATTTGC |
| | | F3 | AGAACGGGAAGCTTGTCATC |
| | | B3 | CGAACATGGGGGCATCAG |
| | | FIP | GACGTACTCAGCGCCAGCATATCTTCCAGGAGCGAGATCC |
| | | BIP | GCGTCTTCACCACCATGGAGACAGAGGGGGCAGAGATGA |

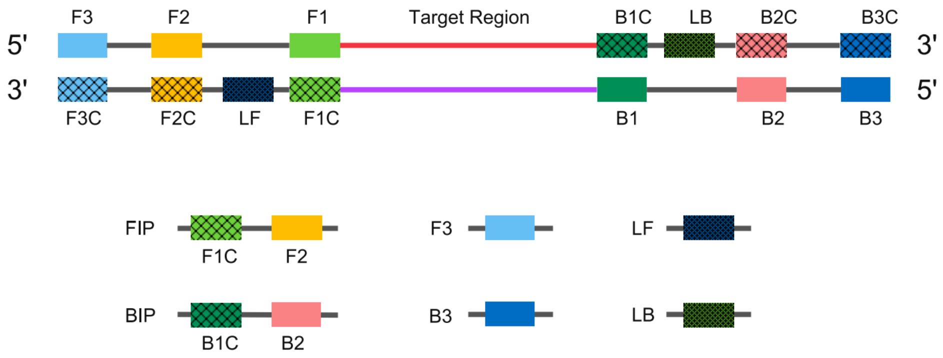


Suppl. Figure 1 LAMP primer structure

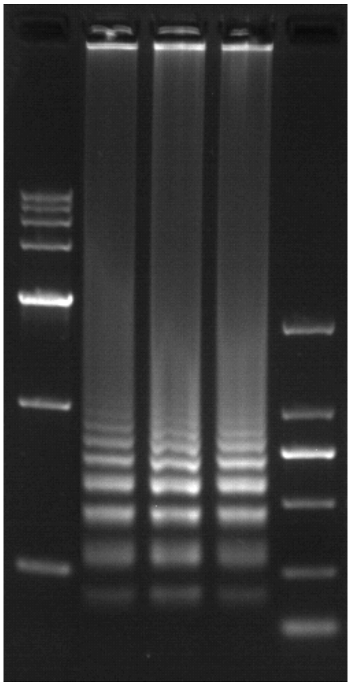

Suppl. Figure 2 Gel electrophoresis image of amplification products

Suppl. Table 2 Nanopore Flongle sequencing data by different time

| Running time | Reads Count | Max Length(bp) | Mean Length(bp) | N50 Length(bp) | Mean Quality |
|---|---|---|---|---|---|
| 30 m | 2263 | 3038 | 440 | 481 | 15 |
| 1 h | 4808 | 3523 | 447 | 488 | 15 |
| 2 h | 6174 | 3523 | 453 | 502 | 15 |
| 12 h | 27000 | 7984 | 483 | 555 | 15 |

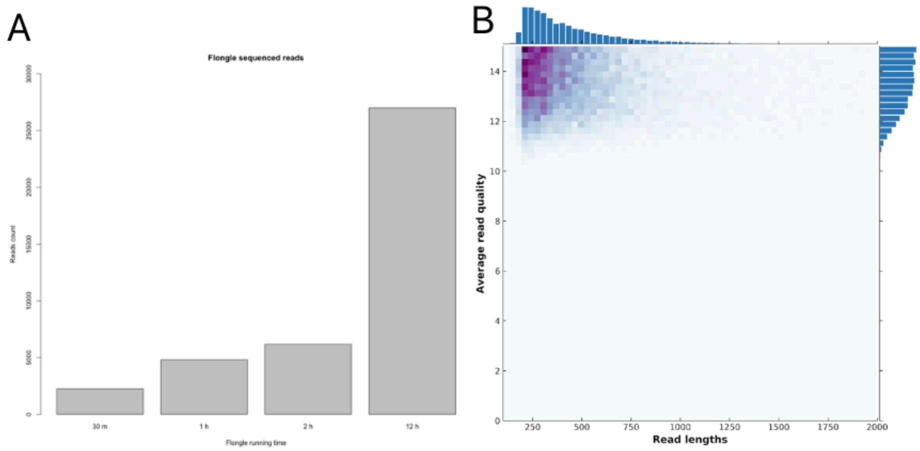


Suppl. Figure 3  Nanopore Flongle sequencing data quality

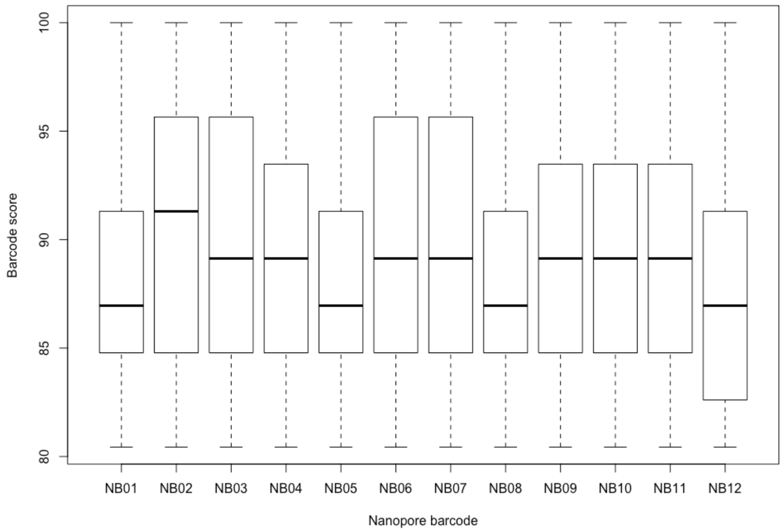


Suppl. Figure 4 Nanopore PromethION demultiplexing barcode score

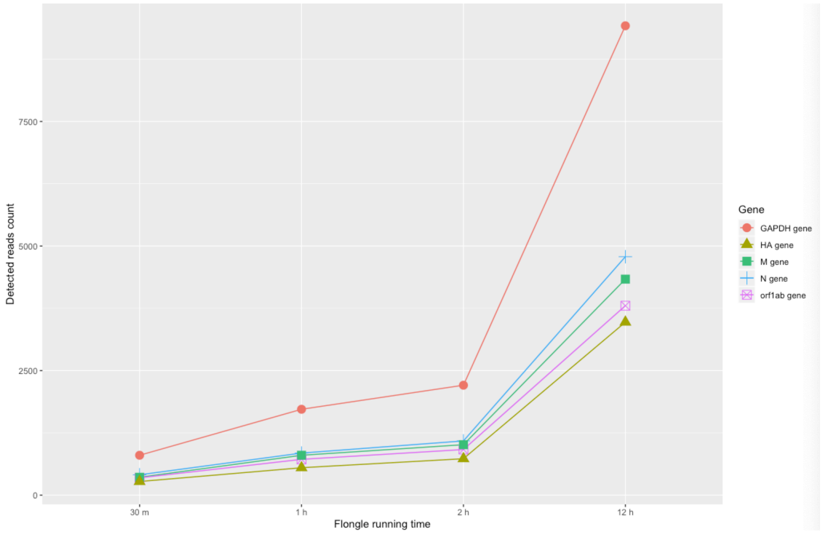


Suppl. Figure 5 Nanopore Flongle detected reads number assigned to target gene

Suppl. Table 3 Nanopore PromethION sequenced and detected reads number

| Sample | Barcode | Sequenced (10^3) | Detected (10^3) | Ratio |
|---|---|---|---|---|
| Positive | NB01 | 217.39 | 187.75 | 0.86 |
| Positive | NB02 | 151.84 | 132.19 | 0.87 |
| Positive | NB03 | 125.48 | 110.46 | 0.88 |
| Positive | NB04 | 21.23 | 17.47 | 0.82 |
| Positive | NB05 | 87.78 | 77.64 | 0.88 |
| Positive | NB06 | 505.27 | 457.08 | 0.90 |
| Positive | NB07 | 429.00 | 385.22 | 0.90 |
| Positive | NB08 | 334.73 | 298.99 | 0.89 |
| Positive | NB09 | 452.65 | 417.21 | 0.92 |
| Positive | NB10 | 135.59 | 121.37 | 0.90 |
| Positive | NB11 | 350.04 | 313.27 | 0.89 |
| Negative | NB12 | 0.21 | 0.01 | 0.07 |
